# Differentially methylated regions and methylation QTLs for teen depression and early puberty in the Fragile Families Child Wellbeing Study

**DOI:** 10.1101/2021.05.20.444959

**Authors:** Roberta De Vito, Isabella N. Grabski, Derek Aguiar, Lisa M Schneper, Archit Verma, Juan Castillo Fernandez, Colter Mitchell, Jordana Bell, Sara McLanahan, Daniel A Notterman, Barbara E Engelhardt

## Abstract

The Fragile Families Child Wellbeing Study (FFCWS) is a longitudinal cohort of ethnically diverse and primarily low socioeconomic status children and their families in the U.S. Here, we analyze DNA methylation data collected from 748 FFCWS participants in two waves of this study, corresponding to participant ages 9 and 15. Our primary goal is to leverage the DNA methylation data from these two time points to study methylation associated with two key traits in adolescent health that are over-represented in these data: Early puberty and teen depression. We first identify differentially methylated regions (DMRs) for depression and early puberty. We then identify DMRs for the interaction effects between these two conditions and age by including interaction terms in our regression models to understand how age-related changes in methylation are influenced by depression or early puberty. Next, we identify methylation quantitative trait loci (meQTLs) using genotype data from the participants. We also identify meQTLs with epistatic effects with depression and early puberty. We find enrichment of our interaction meQTLs with functional categories of the genome that contribute to the heritability of co-morbid complex diseases. We replicate our meQTLs in data from the GoDMC study. This work leverages the important focus of the FFCWS data on disadvantaged children to shed light on the methylation states associated with teen depression and early puberty, and on how genetic regulation of methylation is affected in adolescents with these two conditions.

## 1 Introduction

Epigenetic marks are valuable regulatory mechanisms in health-related studies because they are responsive to the environment and stable across cell division but generally not passed to offspring via meiosis. DNA methylation is a well-established epigenetic marker in which methyl groups are added to the DNA sequence, predominantly at CpG dinucleotides. The CpG dinucleotides often cluster in stretches of DNA termed “CpG islands” [28]. These CpG islands are enriched in the promoter regions of genes, and their methylation can silence or otherwise alter gene transcription [56], and affect many cellular processes, including embryonic development, chromatin structure, X chromosome inactivation, and genomic imprinting [1, 104, 88, 98]. As a consequence, DNA methylation has been found to play an important role in the regulation of many diseases, including cancer [70, 37], neuropsychiatric and neurodegenerative diseases [31], diabetes [75], and cardiovascular disease [74].

Historically, genome-wide association studies (GWAS) have focused on the identification of genomic variation associated with disease [19]. Despite the availability of a complete human genome sequence since 2001 [26, 113, 53], mapping the human methylome has progressed more slowly, mainly due to limitations in technology affecting sensitivity, specificity, throughput, quantitation, and cost. Illumina generated the initial HumanMethylation27 [12] and the more recent HumanMethylation450 (450K) BeadChip [11, 91], which quantifies genome-wide methylation using a cost-effective, high throughput method. These and other advances in methylation quantification now make it possible to study the methylome in greater detail and at a lower cost than ever before.

Because changes in methylation are strongly linked to age [118, 39, 54, 115], these recent technological improvements facilitate studying conditions that occur in specific phases of life. Two conditions of particular interest in adolescent health are teen depression (for all sexes) and early puberty (mainly in girls). These two adolescent conditions both carry higher health risks later in life: Teen depression is associated with an increased risk of obesity [82], anxiety, and suicide [14], and early puberty is associated with an increased risk of breast cancer [4, 106] and asthma [112, 72], and higher BMI [47, 29]. These two adolescent conditions are public health issues, and understanding the regulatory mechanisms governing them would promote the development of preventative treatments and interventions.

Previous work suggests links between methylation and teen depression or early puberty. In particular, several existing studies directly associate differential methylation with depression, but they either do not focus primarily on adolescents [117, 44], are limited to a small sample size [33, 23, 16], examine just a small region rather than the entire genome [51], or only provide data at a single time point [87]. There are also studies identifying differential methylation for early puberty, but again these either have a small sample size [73, 10], only focus on a few candidate genes instead of searching genome-wide [102, 119], or examine individual CpG sites instead of larger methylation regions [22]. Nevertheless, these studies reveal a substantial epigenetic role in both teen depression and early puberty, suggesting that looking for direct associations with methylation could provide valuable insights into the regulatory mechanisms behind both of these conditions.

Here, we aim to further elucidate the role of methylation in teen depression and early puberty. Specifically, we identify differentially methylated regions (DMRs) associated with each of these two conditions, and also DMRs associated with interactions between each of these conditions and age. We then incorporate genotype information to identify genetic variants associated with methylation changes across samples, i.e., methylation quantitative trait loci (meQTLs). Most existing meQTL studies are based on small-sized samples or low-resolution methylation microarrays [41, 124, 8, 94, 6], but nevertheless show promise in identifying novel candidates for explaining mechanisms of disease associations [15, 126]. Finally, we identify interactions of meQTLs with teen depression and early puberty. We thus leverage a relatively large sample of children at two time points to search for meQTLs that can shed further insight into the epigenetic regulation of complex traits, and highlight the effects of teen depression and early puberty on the genetic regulation of methylation.

In this work, our data come from the Fragile Families and Child Wellbeing Study (FFCWS), which is based on a stratified, multistage sample of children born in large U.S. cities (over 200,000 population) between 1998 and 2000 [85]. Births to unmarried parents were oversampled by a ratio of 3 to 1, resulting in a large number of minority and low-income families. Baseline interviews were conducted with mothers in the hospital soon after the child’s birth, and fathers, when available, were interviewed in the hospital or by phone. Follow-up interviews were conducted when the child was one, three, five, nine, and fifteen years old. DNA samples were collected at ages nine and fifteen, and both genome and epigenome-wide methylation assays (Illumina PsychChip arrays version 1.0 and 1.1 for genotypes, and Illumina Infinium HumanMethylation450 BeadChips for methylation) were conducted on subsets of the samples. With these rich data, we identify three types of regulation: i) methylation associated with adolescent depression and early puberty at fixed ages, as well as over time, ii) meQTLs genome-wide, and iii) CpG sites that are genetically regulated differently in adolescents depending on their depression or early puberty status.

Our study adds a unique perspective on the role of methylation in teen depression and early puberty. First, our study examines differences in DNA methylation over time in the same set of participants. Second, our study leverages the longitudinal nature of the data to find methylated regions that are not only associated with teen depression and early puberty, but have interaction effects with age. Third, we are able to use these data to identify large numbers of new meQTLs and also meQTLs with interaction effects for teen depression and early puberty. Thus, our work reveals the complex regulatory relationships between genotype and DNA methylation, and how these relationships change in adolescents with depression and early puberty.

## Materials and Methods

### Data Collection and Processing

#### Study Design and Phenotyping

The Fragile Families and Child Wellbeing Study (FFCWS) identified a birth cohort of unmarried mothers along with a comparison group of married parents in 20 U.S. cities, mainly in the Midwest and on the East Coast. The baseline data were collected between 1998 and 2000, with 4898 mothers interviewed in the hospital within 24 hours of their child’s birth. After this baseline interview, further data were collected from the children at ages 1, 3, 5, 9, and 15, and their mothers. The survey questions at each age span a broad range of important issues in childhood health and well-being; in this work, we focus on teen depression and early puberty as important adolescent traits that also affect future disease risks.

For teen depression, the measured phenotype is a continuous composite score based on multiple questions from the survey data at year 15, with our re-scored scale ranging from 0, indicating no signs of depression, to 15, indicating signs of major depression (Figure 1). This score was adapted from five variables (Table 1) used in the Center for Epidemiologic Studies Depression Scale (CES-D) [64]. These CES-D variables are based on the person’s feelings, with responses on a four-point scale ranging from 1 (strongly agree) to 4 (strongly disagree). In the Fragile Families adaption of the CES-D, the questions pertain to the last four weeks (Table 1). CES-D items were re-scored as follows, to ensure larger scores indicate greater levels of depression and that zero scores are possible: strongly disagree (4 → 0), somewhat disagree (3 → 1), somewhat agree (2 → 2), and strongly agree (1 → 3). “I feel happy” is the only positively worded question; thus, it was re-scored inversely from the other questions, i.e., (4 → 3), (3 → 2), (2 → 1), and (1 → 0). After re-scoring, we generated an overall depression score by summing these items together with all questions weighted equally, following CES-D scoring.

**Figure 1:**
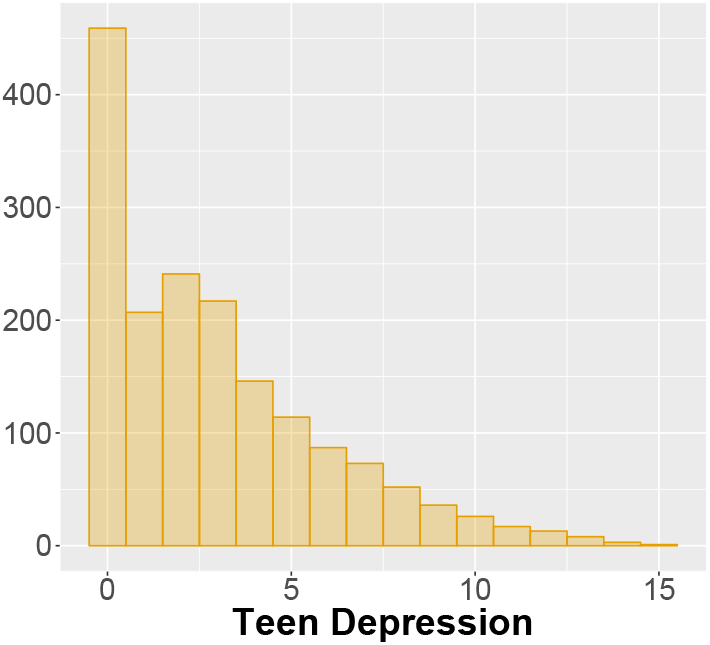
Distribution of teen depression scores in the FFWCS data. The teen depression scores range from 0 (no signs of depression) to 15 (greatest signs of depression). These scores were measured at year 15 across 1700 participants.

**Table 1:** CES-D variables modified for the Fragile Families study in Year 15.

| CES-D Survey Item<br>4-point scale (1-4) | Fragile Families Survey Item |
| --- | --- |
| You felt that you could not shake off the blues, even with help from your family and your friends | I feel I cannot shake off the blues, even with help from my family and my friends |
| You felt sad | I feel sad |
| You were happy | I feel happy |
| You felt life was not worth living | I feel life is not worth living |
| You felt depressed | I feel depressed. |

For early puberty, the measured phenotype is a binary value indicating whether or not breast development is observed at age 9, as recorded in the survey data in the primary caregiver’s response to a question about her daughter, “Would you say that her breasts have started to grow?” We coded responses of “No” as 0, and responses of “Yes, barely” and “Yes, definitely” as 1. This choice was made because any breast development at that age is considered evidence of early puberty [69].

#### Genotype Data and Imputation

The FFCWS collected DNA from saliva samples using DNA Genotek Oragene kits with DNA purification following the manufacturer’s protocol. Genomic DNA was genotyped using Illumina PsychChip arrays versions 1.0 and 1.1.

We performed the following quality control procedure on both PsychChip arrays separately. First, we clustered sample genotypes and removed variant calls that failed quality control (QC) in GenomeStudio. Next, we prepared the genotype data for imputation with IMPUTE2 and 1000 Genomes Project Phase 3 reference haplotype data (October 2014 release) [24, 68]. We aligned genotype alleles to the Human Genome build 37 to match the strands between our genotype data and the 1000 Genomes Project data when possible [84]. The plink algorithms for identical by descent (IBD) detection and sex imputation are sensitive to correlated genetic markers, so we created a version of the data with markers pruned by linkage disequilibrium (LD) using plink `-indep-pairwise 20000 2000 0.5` [81]. We calculated IBD from the LD-pruned data using plink `-genome` and merged a single sample that was duplicated. We checked for erroneous male-female labels by comparing each sample’s reported sex with their imputed sex from X chromosome inbreeding coefficients. We separated the pseudo-autosomal PAR1 and PAR2 regions from the X chromosome and then computed the distribution of F-statistics for males and females based on their observed and theoretical homozygosity. The distribution of F-statistics for the males clustered tightly around 1.0, whereas female F-statistics remained below 0.8. With appropriate sex misclassification thresholds (in plink, `-check-sex 0.8 0.9`), we did not observe sex misclassification in any samples.

In the non-LD pruned data, we removed variants with > 0.05 missingness and individuals with > 0.02 missingness. We discarded erroneous heterozygous calls (heterozygous haploid and non-male Y chromosomes), and we filtered variants (after removing the individuals with substantial missingness above) with a more conservative threshold of > 0.02 missingness. For the PsychChip array version 1.0, the average genotyping rate for the 1212 samples (592 males, 620 females) was 99.9026%; For PsychChip array version 1.1, the average genotyping rate for the 1923 samples (1002 males, 921 females) was 99.8687%. We merged the genotype data from array versions 1.0 and 1.1 using plink (`-merge-mode 1`), resulting in 3119 samples (1586 males, 1533 females), merging genotypes of individuals that were genotyped on both versions of the array. Finally, we ran IMPUTE2 on the 576,200 autosomal variants in 5 Mb windows, yielding 81,694,759 imputed loci (1000 Genomes Project Phase 3 reference, prephasing approach, effective population size 20,000) [68].

#### Methylation Data Collection and Processing

Saliva DNA samples were collected from the focal children at ages 9 and 15 using the Oragene DNA sample collection kit (DNA Genotek). DNA was extracted according to the manufacturer’s protocol and stored at −80 degrees C until use. DNA aliquots (0.5 μg) were bisulfite converted the EZ-96 DNA kit (Zymo Research, Orange, CA) according to the manufacturer’s protocol. DNA was whole-genome amplified, enzymatically fragmented, purified, and applied to the Illumina Infinium HumanMethylation450 BeadChips (Illumina, San Diego, CA) according to the Illumina methylation protocol.

To process the raw methylation data, we used the R package *minfi* [5]. We performed stratified quantile normalization with removal of bad samples as implemented in the *minfi* function `preprocessQuantile`. Next, we removed probes on sex chromosomes and probes with a SNP within one nucleotide of the CpG site, since the presence of SNPs inside the probe body can affect the probe’s binding affinity. We also dropped probes with > 20% failed samples and CpG sites with > 50% failed samples.

Surprisingly, removing probes with known SNPs in them did not fully eliminate SNPs that were perfectly correlated with methylation, indicating a probe with a variant in it rather than a molecular association. Therefore, we tested each site for trimodality, which can indicate the presence of a SNP on the probe body that might have been missed from the SNP filtering step. To do so, for each site, we computed the likelihood ratio test (LRT) statistic comparing a mixture of two Gaussian distributions to a mixture of three Gaussian distributions. These mixture distributions were fit using the R package `mclust` [92]. The LRT statistic was computed as double the difference in log likelihood between the three-component fit and the two-component fit, and was compared to a chi-squared distribution with three degrees of freedom. To set a conservative threshold, we considered a site to be trimodal if its uncorrected p-value was less than 0.05.

To quantify methylation levels, we used M-values, i.e., the logarithm of the ratio between the methylated signal (*M*) and the unmethylated signal (*U*). An alternative quantification of methylation levels is the beta value, which is the ratio between *M* and *M* + *U*. Both M-values and beta values are widely used, but we chose to use the M-value for its improved performance in downstream statistical analyses [34]. After running this pipeline, we were left with a total of *p* = 350,231 CpG sites and *n* = 1444 samples with non-missing sex information to be used in the subsequent analyses.

#### Identification of Differentially Methylated Regions (DMRs)

We examined subsets of the methylation data to identify differentially methylated regions (DMRs) for four variables of interest: *teen depression* (269 samples at age 15 with non-missing measurements), *early puberty* (232 female samples at age 9 with non-missing measurements), *age-depression interactions* (734 samples across both ages with non-missing measurements), and *age-puberty interactions* (363 female samples across both ages with non-missing measurements). Note that there are a larger number of samples in the interaction analyses because many individuals only had complete non-missing measurements for our variables of interest at age 9 or age 15, but not both. For example, if an individual’s early puberty status is known, but their methylation data or BMI value is only known at age 15, they would be included in the age-puberty interaction analysis, where the interaction term would be formed by multiplying 15 by their early puberty status, but not in the puberty analysis because their covariate values at age 9 are missing. For *teen depression* and *early puberty*, the goal is to identify regions of the genome that are differentially methylated across individuals stratified by teen depression levels or early puberty status. For the two age interaction phenotypes, the goal is to identify regions where methylation that changes with age interacts with our variables of interest. In other words, interaction DMRs are genomic regions where the changes in methylation between year 9 and year 15 are themselves different between individuals stratified by their depression score at age 15 or their early puberty status at age 9.

To identify DMRs and interaction DMRs, we first identified associations between methylation at single CpG sites and the variable of interest (teen depression, early puberty, age-depression interactions, or age-puberty interactions). These site-level results were then combined locally across the genome to identify differentially methylated regions.

#### Associations with Single CpG Sites

To identify the methylation of single CpG site-level associations with our phenotypes of interest, we followed a two-step procedure. First, we performed principal component analysis (PCA), once on the standardized methylation data and once on the imputed standardized genotype data. We selected the first two principal components (PCs) from both matrices to use in association testing to control for batch effects and population structure [63, 80]. The methylation PCs are computed separately for each analysis (depression, early puberty, age-depression interaction, and age-puberty interaction) by subsetting to include only the subjects with non-missing covariate information for that association test.

Second, we fit linear regression models to the M-values of each CpG site. We define the M-values of a CpG site as *Y_i,j_*, where *i* = 1,…, *n* is the sample index and *j* = 1,…, *p* is the CpG site index. Let *Z_i_* be the relevant covariates for a model, and *X_i_* be one of the four variables of interest (i.e., the measure of teen depression, early puberty, age-depression interaction, or age-puberty interaction). Then, we fit the model

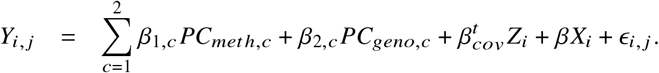

For each of the four variables of interest, different covariates were included in the model. For 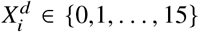 representing teen depression status, we used the continuous score described above (higher values indicate more extreme depression), with the covariate sex, which is encoded as a binary value. Early puberty 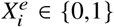 is a binary value where a 1 indicates breast development has started at age 9, and the covariate is the real-valued BMI measured at age 9. For age-depression interactions, we examined the interaction term between age—a binary variable indicating whether the sample is from year 9 or year 15—and the depression score at age 15. The covariates were sex, age, and depression score. Finally, for age-puberty interactions, we examined the interaction term between age and early puberty status, and the covariates were age, early puberty status, and BMI. The interaction terms were defined as the product of the two numeric values for each sample (binary values are made numeric with {0,1} dummy variable encoding). All models were fit for each CpG site using the R package *limma* [99]. The built-in empirical Bayes function was used to shrink the standard errors towards a common value, and p-values were reported for each variable of interest and each CpG site from these model fits.

#### Identifying Regions of Interest from Summary Statistics across Single CpG Sites

We are primarily interested in differentially methylated regions (DMRs), rather than considering each CpG site marginally, due to the greater statistical reliability of differential methylation across regions. Thus, we combined the individual p-values for each CpG site across genomic regions to identify differentially methylated regions. To do so, we applied a “moving averages” method of p-value correction [79] that accounts for spatial correlation and multiple hypothesis test correction to identify DMRs from the p-values of each CpG site. We first computed the correlation between CpG sites at a varying genomic distance (i.e., autocorrelation). Second, we performed the Stouffer-Liptak-Kechris correction [57], where each p-value is modified with a weight derived from the autocorrelation of the associated CpG site with other CpG sites within the genomic window. Third, we used the Benjamini-Hochberg correction [9] to control the false discovery rate (FDR). The resulting q-values for each CpG site were used to identify DMRs based on the seed and distance parameters. We set the seed to *q* ≤ 0.2 (maximum q-value required to start a region), and the distance to 1000 bases (distance in which another sufficiently low q-value must be found for the genomic window to be extended). Finally, we computed the Stouffer-Liptak p-value for each region by using the autocorrelation calculated out to the length of the longest region, and we applied the Sidak correction [96] for multiple testing on the Stouffer-Liptak p-values for each region. These Sidak-corrected p-values are reported for each potential differentially methylated region. All these steps were performed using the Python package `comb-p` [79].

#### Methylation Quantitative Trait Loci (meQTLs)

We identified three types of methylation quantitative trait loci (meQTLs), described below. The baseline meQTLs were ascertained from the full sample (*n* = 748) by testing for association between CpG sites and genotypes. The two types of interaction meQTLs were ascertained by testing for an interaction that includes the single nucleotide polymorphism (SNP) and teen depression or early puberty—the effect of the association between SNP and CpG site changes as a function of teen depression or early puberty status—while also including both the SNP and the covariate of interest as fixed effects in the model.

#### Methylation QTLs

To identify meQTLs, we tested each CpG site and SNP pair for associations between methylation levels and genotype using *MatrixEQTL* [93]; although the package was designed for expression QTLs, we considered M-values of each CpG site across samples instead of transcript expression levels. This package is often used for identifying meQTLs [30, 67, 77]. We corrected for multiple comparisons by computing the false discovery rate (FDR) using Benjamini-Hochberg [9] and using an FDR < 0.05 threshold for discovery. In our analysis, we examined only cis-meQTLs (distance of the SNP from the CpG site ≤ 20 kb) using all 748 samples with non-missing covariate information [6], and we fit the following linear regression model for each SNP, CpG site pair:

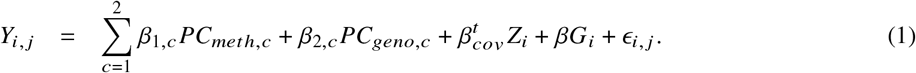

Here, *G_i,l_* ∈ {0,1,2} is the genotype for individual *i* at position *l* ∈ {1,…, *L*}, encoded as the number of copies of the minor allele. In this analysis, we included as covariates *Z_i_* sex and age, and two PCs each for methylation and genotype as in the DMR analyses [6].

#### Interaction meQTLs

We used this same association testing approach to find *interaction meQTLs,* or meQTLs that have interaction effects with teen depression and early puberty, by including the appropriate interaction term in the model above (Eqn 1). In other words, we found meQTLs where the association between the CpG site and SNP is differential with respect to either the depression score or early puberty status. All samples with a depression measurement at year 15 were used to find the teen depression interaction meQTLs, controlling for sex, age, and depression score at year 15. The female year 9 samples were used for the early puberty interaction meQTLs, controlling for BMI and early puberty status. The four PCs (two from methylation measurements and two from genotypes) were also included as covariates in all models.

#### meQTL Validation Analysis

The Genetics of DNA Methylation Consortium (GoDMC) [71] focuses on epidemiological studies with more more than 30,000 participants with genetic, phenotypic and DNA methylation information. In particular, the GoDMC project identified meQTLs from whole blood DNA samples using HumanMethylation450 or EPIC array profiles from 32,851 participants in the GoDMC UK cohort [71]. We validated our meQTL results by comparing them to the GoDMC study’s meQTLs. In particular, we examined the GoDMC meQTL discoveries in the phase 2 analyses, which consists of all significant and non-significant SNP-CpG results in the GoDMC data, in light of the meQTL associations we identified in the FFCWS data.

We performed the validation as follows. First, using the meQTLs from GoDMC, we identified the same CpG-SNP pairs in our study (85,984,217 GoDMC meQTLs; 9,154,359 meQTLs in our study; 48,566 unique SNP and CpG pairs in common). The corresponding p-values of the two studies, FFCWS and GoDMC, were plotted in a scatterplot to check their trend. Finally, we also compared the density plot of the p-values found in the FFCWS study data sets and the p-values found in the GoDMC data.

#### meQTLs Enriched in Cis-Regulatory Elements

We identified enriched genomic functional elements using stratified linkage disequilibrium (LD) score regression [38]. This method identifies meQTLs that are enriched in specific genomic elements by comparing the genetic contribution to the variance of the methylation values of groups of SNPs in LD that are colocalized inside and outside of the functional element. We used the Python *ldsc* package [18] to perform enrichment tests for 53 overlapping functional element categories from 24 top-level annotations, using the default base-pair windows and the 1000 Genomes Project European-ancestry GWAS data LD Score regression intercepts. The 24 top-level annotations considered are: UTR; coding; regions conserved in mammals [66, 116]; open chromatin reflected by DNase I hypersensitivity site (DHS) regions [111, 45]; superenhancers [49]; FANTOM5 enhancers [3]; histone marks H3K4me1, H3K4me3, H3K9ac [36, 60, 111]; introns, promoter regions [45, 58]; and the combined chromHMM and Segway annotations [50].

## Results and Discussion

Our results include two primary analyses: DMRs and meQTLs. The goal is to discover both methylation levels associated with teen depression and early puberty, and how genotypes regulate methylation levels differently in participants with the two conditions.

### DMRs for Teen Depression

First, we identified genomic regions that are differentially methylated across Fragile Families participants with and without teen depression. With 269 participants, we detected 4 DMRs (corrected p-value ≤ 0.05; Table 2a). To understand the implications of these results and the extent to which our findings agree with previous results in the literature, we examined the genes overlapping each of the DMRs. We found one gene overlapping these four regions, *ZSCAN1*. Although this gene has not been found directly linked to depression, *ZSCAN1* is involved in high-risk HPV, inducing carcinogenesis [114], and in cervical neoplasia [35]. Our findings suggest new insights for genes with previous associations with other diseases.

**Table 2:** Summary of the four (corrected p-value *p* ≤ 0.05) DMRs (a) associated with teen depression, and a summary of the five (corrected p-values *p* ≤ 0.05) DMRs (b) associated with age-depression interaction. Genes overlapping any of these regions are reported in the *Gene* column. The *p* column represents corrected *p*-values.

| (a) |  |  |  |  |  | (b) |  |  |  |  |  |
| --- | --- | --- | --- | --- | --- | --- | --- | --- | --- | --- | --- |
|  | Chr | Start | Stop | Gene | *p* |  | Chr | Start | Stop | Gene | *p* |
| 1. | 8 | 37824496 | 37824766 | - | 0.0006302 | 1. | 4 | 165898707 | 165898968 | *TRIM61* | $1.75 \times 10^{-5}$ |
| 2. | 6 | 28446794 | 28447116 | - | 0.006125 | 2. | 13 | 19918951 | 19919209 | - | 0.0031 |
| 3. | 19 | 58554401 | 58554528 | *ZSCAN1* | 0.0225 | 3. | 19 | 7852051 | 7852239 | *EVI5L* | $5.83 \times 10^{-6}$ |
| 4. | 19 | 7852051 | 7852239 | - | $2.391 \times 10^{-5}$ | 4. | 11 | 1474456 | 1474778 | *BRSK2* | 0.0086 |
|  |  |  |  |  |  | 5. | 8 | 11203882 | 11204133 | - | 0.00197 |

### DMRs for Early Puberty

Next, we identified genomic regions that are differentially methylated across those with and without early puberty status. We considered 232 participants, 128 of whom were positive for early puberty, and we found ten DMRs (corrected p-value *p* ≤ 0.05; Table 3a). Of the six genes overlapping these DMRs, we highlight two genes of interest for the early puberty phenotype: *ABAT* and *GRK4*.

**Table 3:**
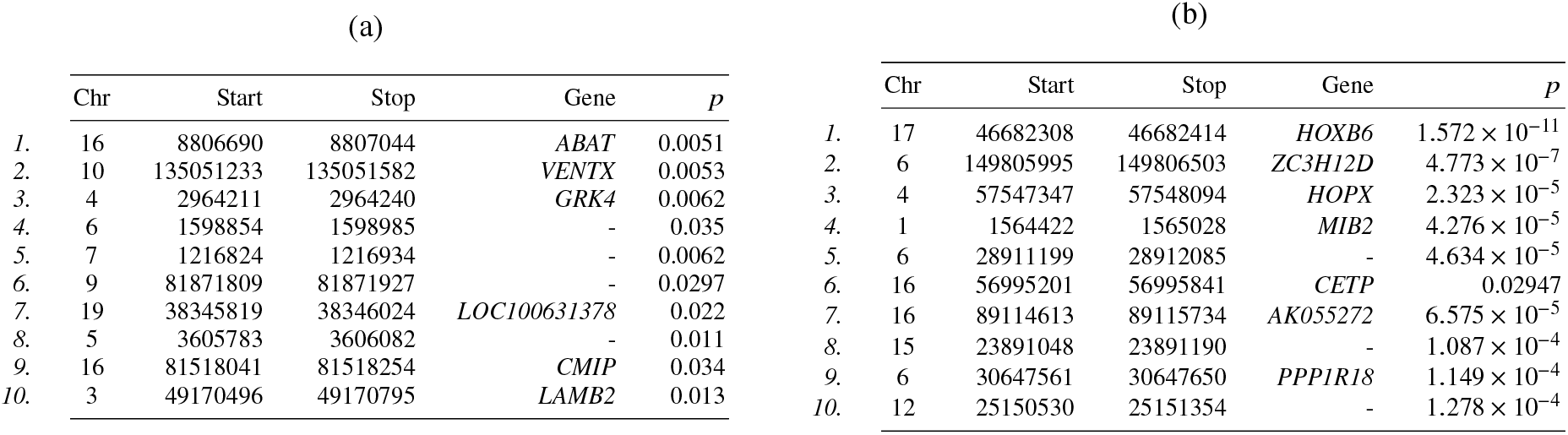
Summary of the ten differentially methylated regions (DMRs; corrected p-value *p* ≤ 0.05) associated with early puberty (a), and the top ten DMRs (out of 80) associated with age-puberty interactions (b). Genes overlapping these regions are reported in the *Gene* column. The *p* column represents corrected *p*-values.

*ABAT* encodes a mitrochondrial enzyme that is a key regulator in the metabolism of a major inhibitory neurotransmitter known as gamma-aminobutyric acid (GABA). GABA plays a major role in a wide range of brain functions, and its dysfunction due to mutations in *ABAT* has been linked to many different disorders, including epilepsy, psychomotor difficulties, and Alzheimer’s disease [127]. A study on puberty in goats found *ABAT* to be one of the implicated genes in methylation changes before and after puberty [121], and this gene was also found to be important in puberty and reproductive functions in rats [95].

Next, *GRK4* encodes the G protein-coupled receptor kinase 4. While no direct association has been previously observed with early puberty, this gene has been found to be associated with BMI in adolescent girls [90]. Childhood BMI is known to be associated with pubertal timing [40], which could suggest a link between *GRK4* and early puberty. Moreover, G protein-coupled receptor kinase 4 is related to hypertension and lipid levels [100]. Correlations between lipid levels and hypertension with BMI in prepubertal children have been established [17]. Thus, *GRK4* appears to be involved in early puberty through childhood BMI, lipid levels, and hypertension.

We did not find relevant annotations for the remaining four genes. Taken together, our DMRs have biological support since the overlapping genes are enriched for plausible connections with early puberty, while also suggesting possible mechanisms for early puberty and highlighting other potentially mechanistic genes.

### DMRs for Age-Depression Interaction

We next tested for DMRs associated with an age-depression interaction, meaning that the effect size of a DMR for depression is associated with age. We found five age-depression interaction DMRs (corrected p-value *p* ≤ 0.05; Table 2b; Figure 2A).

**Figure 2:**
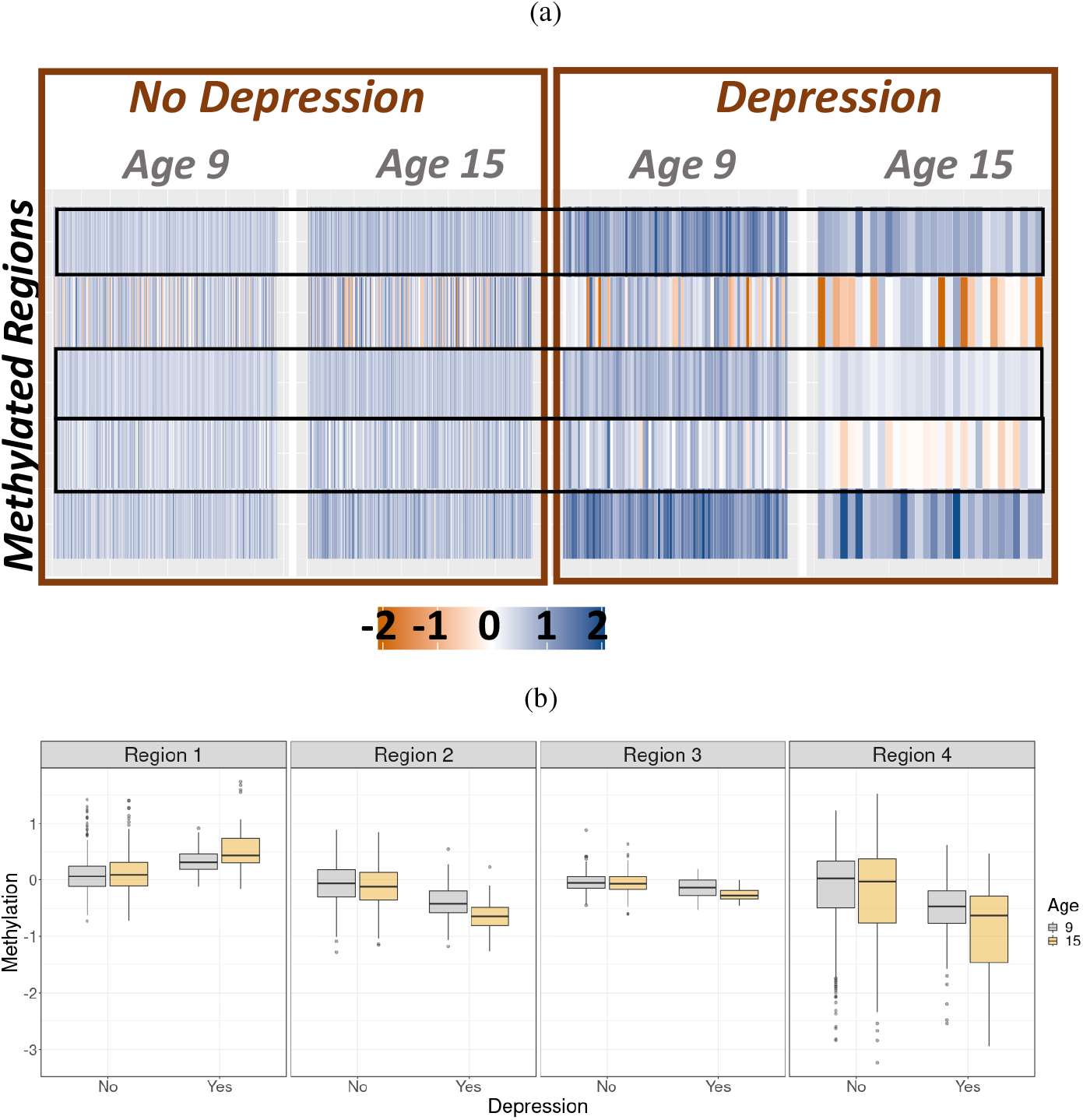
Interactions between age and depression for differential methylation. Residualized M-values are computed by removing the effects of the associated covariates that are not the phenotypes of interest. For visual purposes, we stratify the depression scores by those less than or equal to six versus those greater than six. (a) Heatmap of the methylation levels in DMRs for age-depression interactions (Table 2b). Methylation levels are quantified as residualized M-values. (b) Boxplots of the four highlighted DMRs (numbers 1, 5, 10, and 14 in Table 2b) as examples of differential methylation at these regions between ages 9 and 15 among one of the categories (depression or no depression) with no evidence of differential methylation among the other category. These regions show an interaction effect between age and depression on methylation.

We found three genes of interest overlapping our age-depression DMRs: *TRIM61, EVI5L,* and *BRSK2* (Figure 2B). Of these, *BRSK2* has a previously studied association with disorders of the brain. In particular, mutations in this gene have been found to be related to neurodevelopmental disorder [48]. A separate study identified the BRSK2 kinase to suppress a transcription factor known as NRF2, which is relevant because NRF2 activity is known to decrease in neurodegenerative disorders as well as with age [105]. Hence, our finding suggests that this gene could have an effect on depression that is modified by age.

### DMRs for Age-Puberty Interactions

Next, we tested for associations among methylation levels at each CpG site and the age-puberty interaction term, which tests whether the effect sizes of early puberty DMRs are a function of age. We identified 80 age-puberty interaction DMRs (corrected p-value ≤ 0.05; Table 3b). Interestingly, many more age-puberty DMRs were found relative to the number of early puberty DMRs detected. This suggests that early puberty may primarily manifest in methylation markers dynamically across ages rather than as differences detectable at a fixed age. Across the 10 most significant DMRs (reported in Table 3b), the identified age-puberty effects appear to have no differential effects in the controls (children without early puberty), and have the same direction of differential effect in the age 9 and age 15 cases (children with early puberty). For the majority of our interactions, the magnitude of the differential effect is larger for the age 15 cases than for the age 9 cases (Figure 3a).

**Figure 3:**
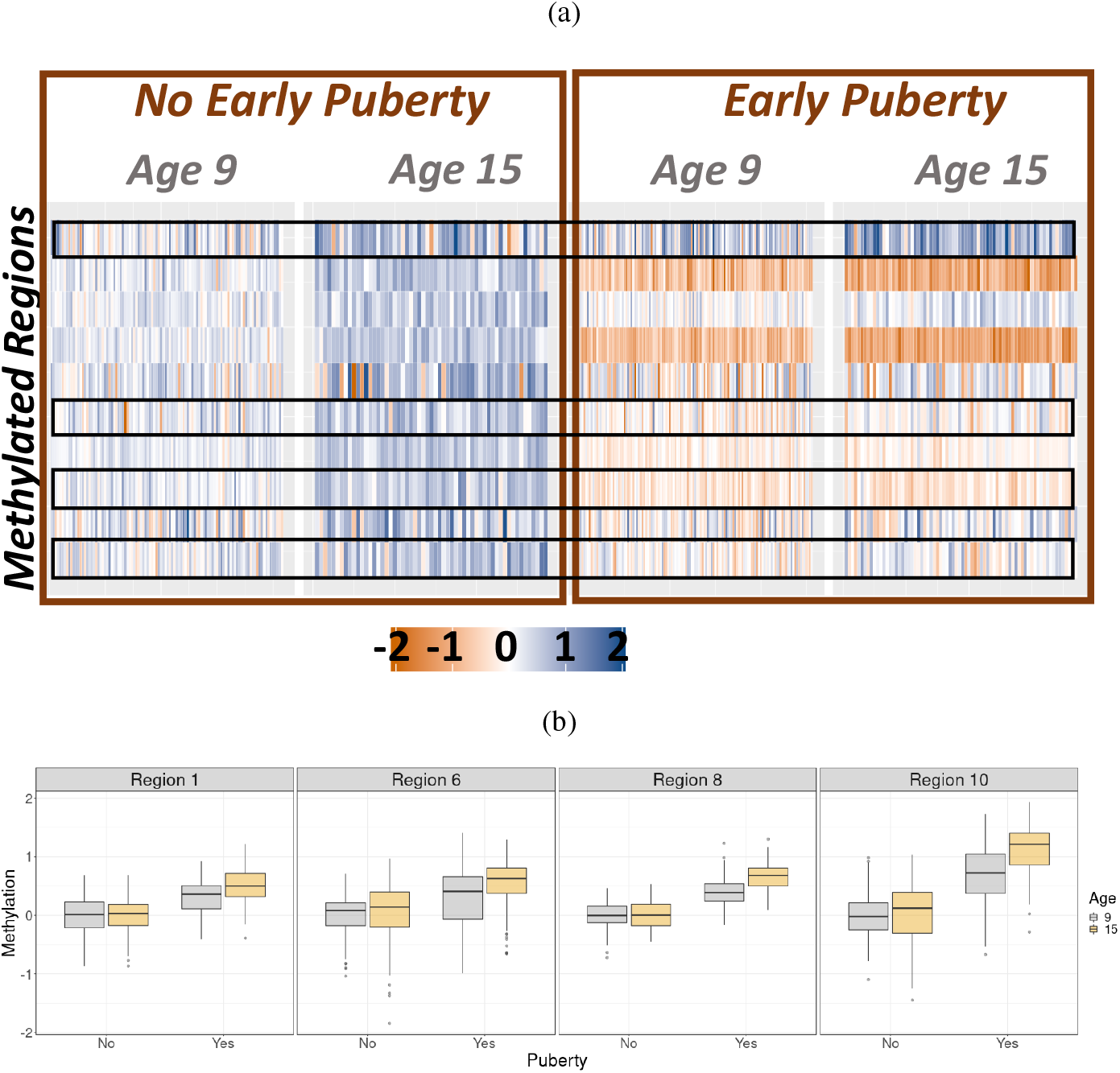
Interactions between age and early puberty associated with differentially methylated regions. Residualized M-values are found by removing the β coefficient with the associated covariates that are not the phenotype of interest. (a) Heatmap of methylation levels in the DMRs for age-puberty interaction terms (Table 3b). Methylation levels here are measured as residualized M-values. (b) Boxplots of four DMRs (1, 5, 10, and 14 in Table 3b) as examples of differential methylation at these regions between ages 9 and 15 among one of the categories (early puberty or no early puberty), but no differential methylation between the two ages among the other category.

To study the functional impact of these interaction DMRs, we examined the seven genes overlapping the top ten associated regions; here, we highlight one of them (CETP), and also the regions overlapping CETP to show the differences in effects (Figure 3b). *CETP* is known to have associations with HDL cholesterol levels, and mutations at this locus can result in greater risk of myocardial infarction [86]. This gene is also specifically shown to have associations with lipid levels in both children [97]. This suggests particular relevancy here, since BMI is known to be related to pubertal timing, and lipid levels are associated with BMI.

Overall, the DMRs we found associated with the four variables of interest are important for three reasons. First, we are able to validate and strengthen previous results concerning particular genes and their associations with depression and early puberty. Second, we provide a possible mechanism of action for these variables of interest, by associating these genes with the variables of interest through increased or decreased methylation. Our interaction DMRs in particular highlight genetic loci whose methylation is differential by age conditional on teen depression or early puberty status, suggesting a dynamic, rather than static, association and thus greater context for these mechanisms across relevant ages. Third, our findings generate novel insights by implicating genes that have not been directly connected to these specific variables of interest. Moreover, the remaining regions that do not directly overlap genes could still represent important findings about non-coding regions that may play a functional role in the regulation of teen depression and early puberty.

### Methylation QTLs

We shift our focus to the second methylation phenomenon of interest: Methylation quantitative trait loci (meQTLs), or genetic variants that are associated with CpG site methylation levels. First, we describe a genome-wide analysis of the full sample (*n* = 748) to identify associations between all CpG sites and single nucleotide polymorphisms (SNPs) in our FFCWS data. We then perform enrichment analysis to further characterize possible mechanisms of these meQTLs. Second, we show meQTLs that have interaction effects with depression or early puberty and again perform enrichment analysis on these interacting meQTLs. Last, we validate the puberty-interacting meQTLs in the GoDMC data set to demonstrate the generalizability of our findings [76].

### Genome-Wide Methylation QTLs

We first identified meQTLs across 748 samples. Our analysis discovered 6,103,375 SNP-CpG pairs representing meQTLs (FDR ≤ 0.05; Tables 4), including a total of 2,835,920 unique SNPs and 209,624 unique CpG sites. Many SNPs and CpG sites were implicated in multiple associations, highlighting the correlations among CpG sites nearby on the chromosome with shared meQTLs, as with SNPs in LD that were associated with the same CpG site (Figure 4). This result is obtained by removing trimodal CpG sites from the analysis with the likelihood ratio test procedure; the trimodal distribution identified CpG sites that were often perfectly correlated with a nearby SNP, indicating that the probe may have been affected by the SNP instead of effects happening at a cellular level. We did not test for associations between non-local SNPs and CpG sites, so distal sharing cannot be observed in our results; however, the local shared effects are also reflective of the local correlation structure known for CpG sites [125] and for SNPs via linkage disequilibrium.

**Table 4:** Fifteen meQTLs at unique genetic loci with lowest p-values, with the corresponding effect size (*β* value). If a CpG site is located within a gene, the gene name is indicated in the *Gene* column. The *p* column represents corrected *p*-values.

| Chr | SNP Position | CpG Position | Gene | Beta | $p$ |
| --- | --- | --- | --- | --- | --- |
| 1 | 161360435 | 161349006 | *SDHC* | 1.81 | $2.22 \times 10^{-308}$ |
| 6 | 41196605 | 41195891 | *TREML2* | -1.26 | $2.22 \times 10^{-308}$ |
| 2 | 30670010 | 30669759 | - | -1.74 | $2.22 \times 10^{-308}$ |
| 11 | 64115983 | 64116279 | *MACROD1* | 1.72 | $2.22 \times 10^{-308}$ |
| 15 | 41689232 | 41695430 | - | 1.71 | $2.22 \times 10^{-308}$ |
| 17 | 75961513 | 75956447 | *ACOX1* | 1.49 | $2.22 \times 10^{-308}$ |
| 4 | 122720999 | 122721982 | - | 1.63 | $2.22 \times 10^{-308}$ |
| 10 | 134150920 | 134165390 | - | 2.12 | $2.22 \times 10^{-308}$ |
| 1 | 51811558 | 51811558 | - | 3.40 | $2.22 \times 10^{-308}$ |
| 8 | 144699424 | 144715010 | - | -1.73 | $2.22 \times 10^{-308}$ |
| 12 | 120537408 | 120535099 | - | 1.81 | $2.22 \times 10^{-308}$ |
| 16 | 22166976 | 22175965 | - | 4.56 | $2.22 \times 10^{-308}$ |
| 7 | 45025696 | 45025080 | *CCMD2* | 1.73 | $2.22 \times 10^{-308}$ |
| 6 | 28881660 | 28885017 | - | 1.21 | $5.10 \times 10^{-303}$ |
| 19 | 39953823 | 39967363 | - | 1.92 | $1.94 \times 10^{-304}$ |

**Figure 4:**
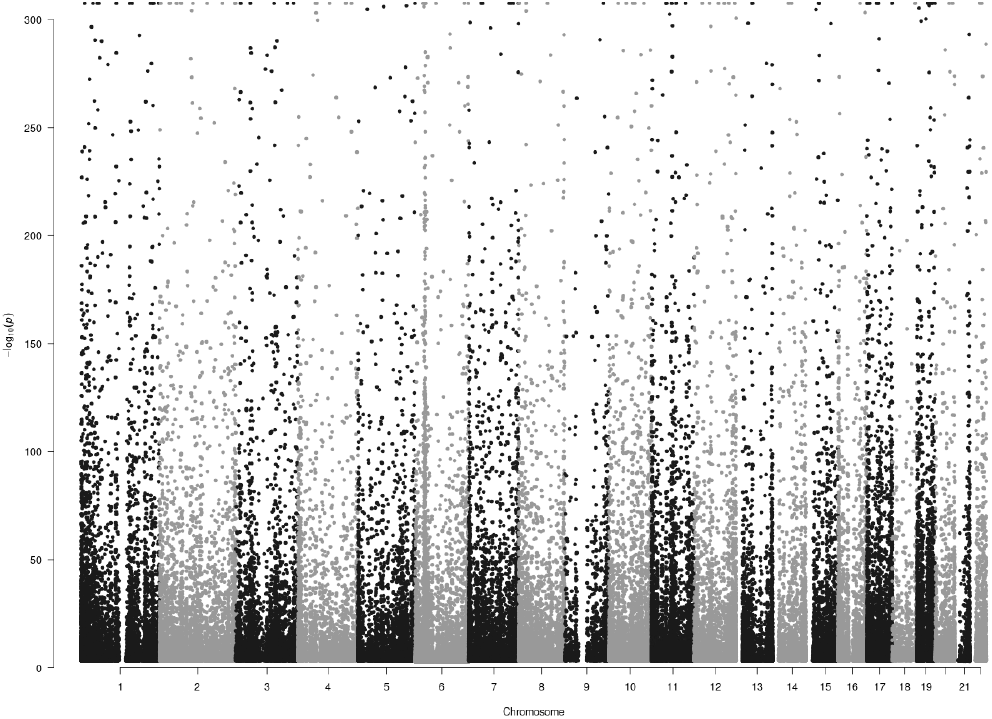
Manhattan plot for each SNP showing the -log_10_ p-value with each associated CpG site, for all cis-meQTLs with p < 1 × 10^−3^.

We further identified several genes of interest overlapping CpG sites in the top 15 meQTLs in unique genomic loci. In particular, the *TREML2* gene is associated with Alzheimer’s disease and other neurological disorders [120].. Next, the *NRG3* gene has been found to be related to cognitive deficit in schizophrenia [65], as well as bipolar disorder and major depressive disorder [78]. Finally, mutations in the *ACOX1* gene are associated with neurodegeneration [42]. All three genes play important roles in biological processes, and impact psychiatric and general health in a variety of ways. Our meQTL findings contribute by showing that colocalized CpG sites are regulated by genotype, suggesting a specific genetic regulatory mechanism at play.

Next, we performed enrichment analysis using stratified LD score regression on the meQTLs to quantify enrichment of genetic effects in functional annotations [38]. Each hypothesis test evaluates whether the meQTLs are enriched in one of the 24 annotation categories or not (Bonferroni-corrected *p* ≤ 0.05; Figure 5). We observed enrichment for 18 of these annotation categories. A few of these categories are of particular interest in mechanistically understanding meQTLs. First, FANTOM5 enhancer regions [3] showed the greatest enrichment among all annotations with 0.4% of the SNPs explaining 14.8% of meQTL heritability. Enhancers are thought to functionally regulate gene expression more distally than promoters [3]. Enhancers have been found to have a direct relationship with early puberty [108, 109], where the repression or activation of enhancer regions is essential for regulating the timing of mammalian puberty. Super-enhancers did not show a large enrichment compared to enhancers; the enrichment estimate is 2.2 for super-enhancers compared to 14.8 for the FANTOM5 enhancers. Weak enhancers, on the other hand, showed substantial enrichment, with 2.1% of the SNPs explaining 47.03% of the meQTL heritability. Although still an enrichment, the smaller estimate for super-enhancers suggests that they may not play as big a role in meQTLs compared to other enhancer regions.

**Figure 5:**
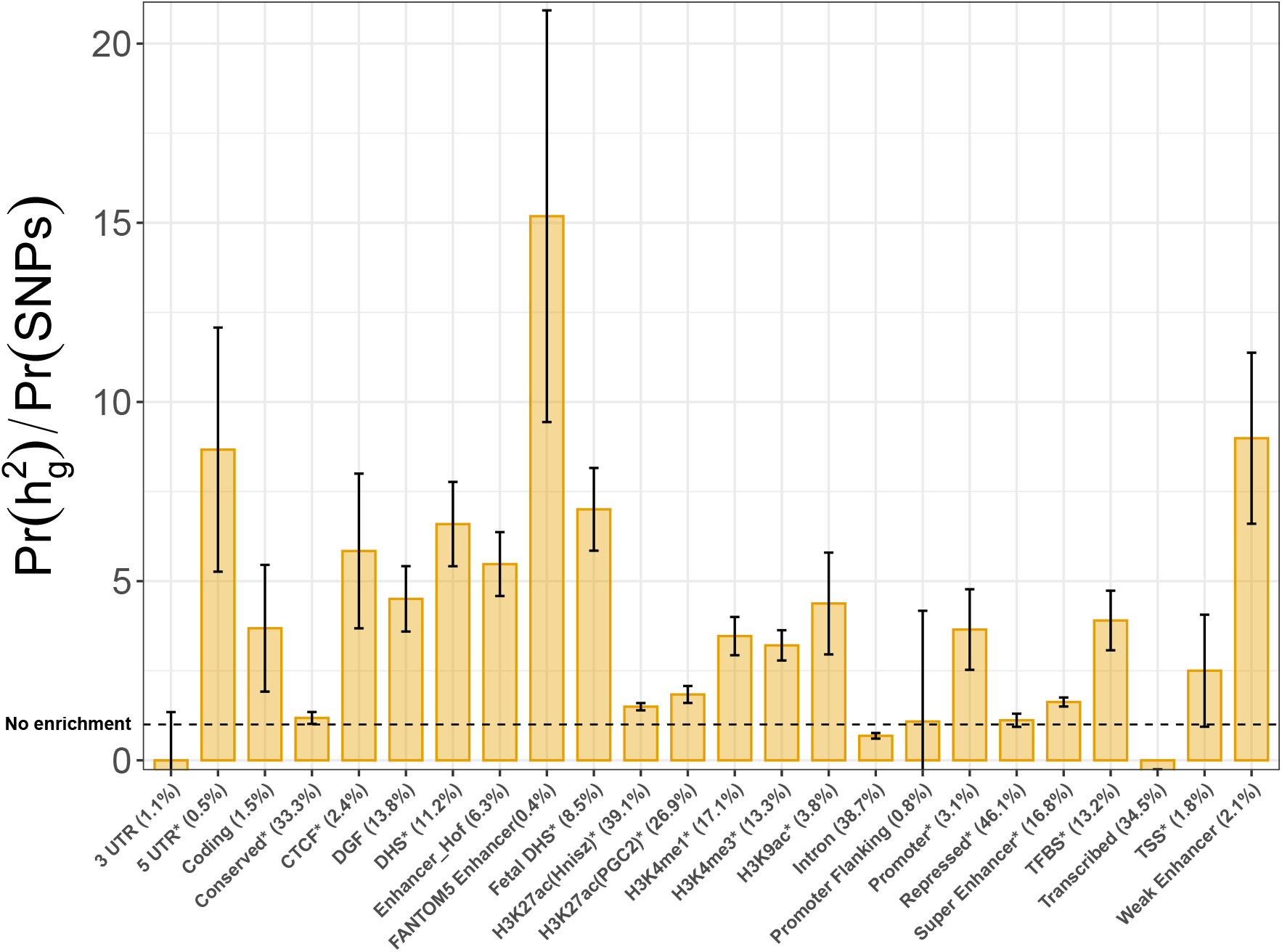
Enrichment analysis using stratified LD score regression for the 24 regulatory element categories. Error bars show the jackknife standard errors around the estimates of enrichment, with the asterisk representing significance (p ≤ 0.05) after Bonferroni correction for the 24 hypotheses tested. The percentage in parentheses indicates the percentage of SNPs colocalizing with the regulatory element.

Second, the 5’ untranslated region (5 UTR) also shows substantial enrichment, with 0.5% of the SNPs explaining 11.4% of the meQTL heritability. The *5* UTR has been found to be related to depression [89], schizophrenia [61], and attention-deficit hyperactivity disorder [7].

### Validation in the GoDMC Study

The GoDMC study identified meQTLs calculated in whole blood DNA samples with HumanMethylation450 or EPIC array profiles from 32,851 participants in the GoDMC cohort [71]. We compared our meQTLs with the meQTLs identified in the GoDMC study to validate our results out-of-sample. Among the 7,786,541 CpG-SNP pairs in common between the two studies (with 348 unique SNP and CpG pairs in common), we see a strong enrichment of p-values found in the FF meQTLs as compared to the p-values obtained in the GoDMC data (Figure 6; Section §B.1). This validation shows that the meQTLs identified in the FFCWS replicate in an outside sample, and suggests that the meQTLs represent reproducible biological signal.

**Figure 6:**
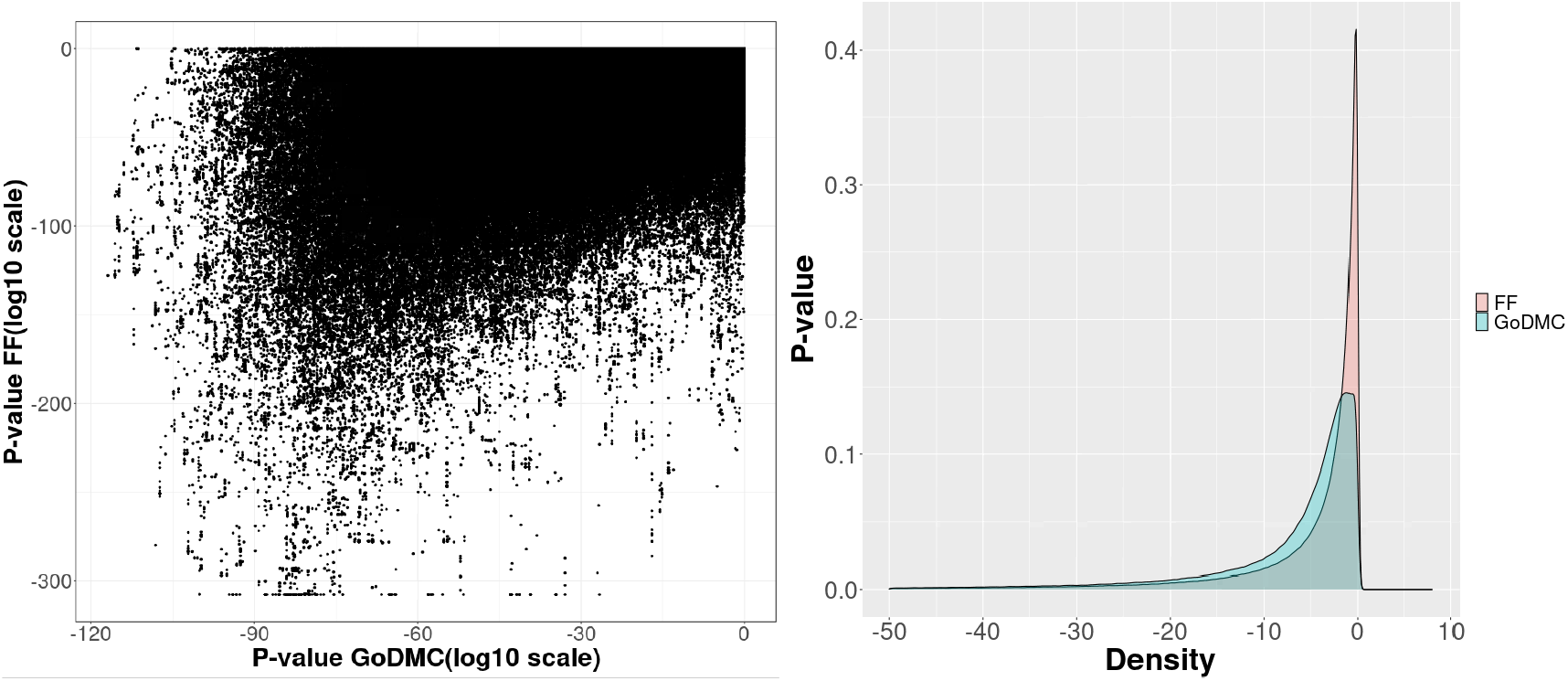
FFCWS meQTL replication in the GoDMC Study. (a) Scatterplot of the log10 p-values found in the FFCWS data versus the log10 p-values found in the GoDMC data from CpG-SNP pairs in common between the two studies. (b) Density plot of the p-values found in the FFCWS data on log10 scale (pink) and the p-values found in the GoDMC data (in blue) from CpG-SNP pairs in common between the two studies.

### Interaction meQTLs

Next, we looked at interaction meQTLs to find SNP-CpG site associations that are differential based on early puberty or teen depression status. To do this, we included the appropriate interaction term in the linear models for each pair of SNPs and CpG sites (see § Methods). We found a total of 13,658 interaction meQTLs for puberty, involving 13,269 unique SNPs and 4,532 unique CpG sites. We found 21,543 interaction meQTLs for teen depression, involving 20,943 unique SNPs and 8,525 unique CpG sites (FDR ≤ 0.05; Table 5).

**Table 5:**
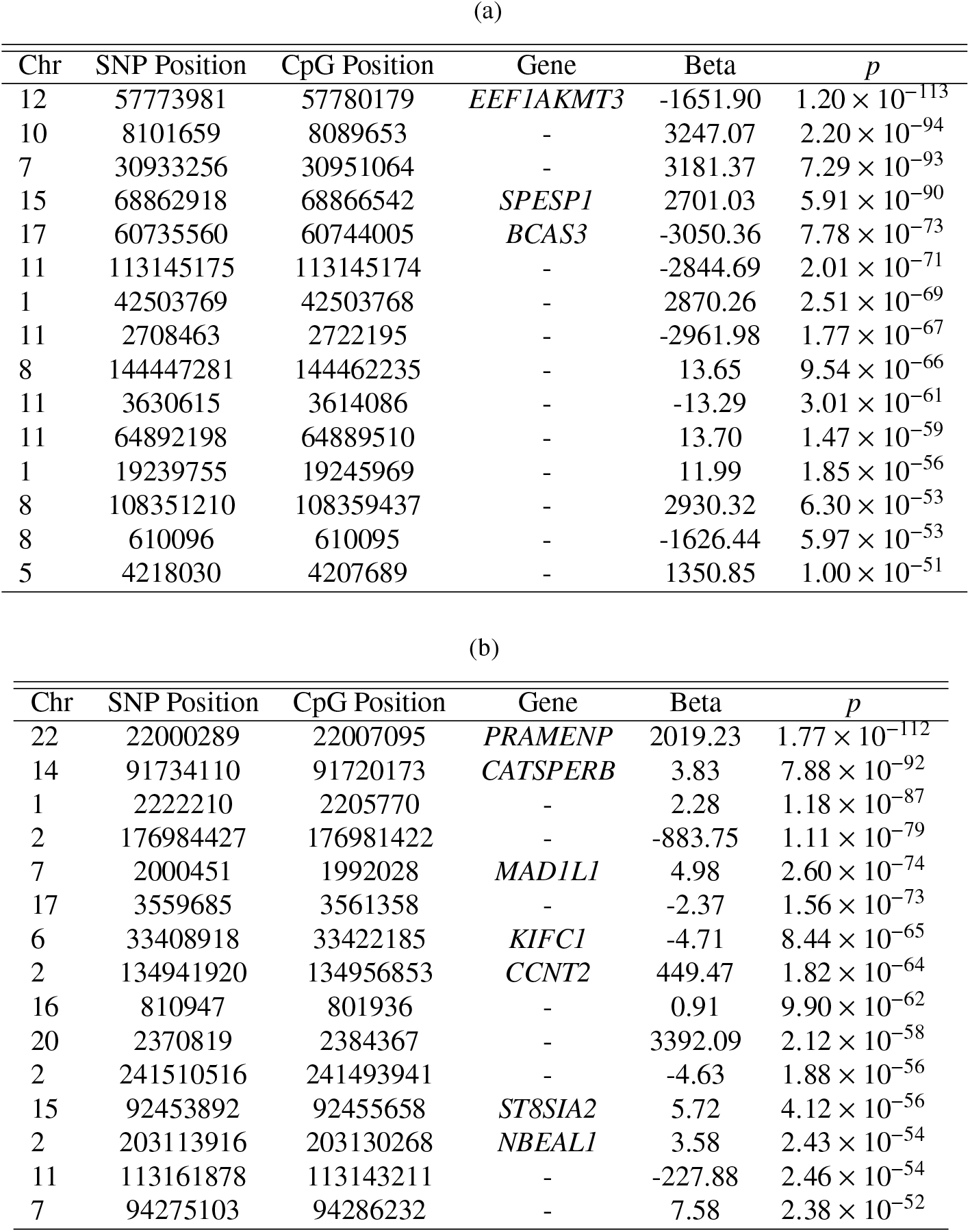
Puberty and depression interaction meQTLs. (a) Fifteen puberty interaction meQTLs with the lowest p-values with the corresponding estimated effect sizes (*β* values). If a CpG site is contained within a gene, the gene is indicated in the *Gene* column; (b) Fifteen depression interaction meQTLs with the lowest p-values, with the corresponding estimated effect size (*β* values). If a CpG site is contained within a gene, the gene is indicated in the Gene column. The *p* column represents corrected *p*-values.

To illustrate the interaction effects, we examine a few meQTLs with depression or early puberty interactions more closely (Figure 7). For the teen depression interaction meQTLs, the association between SNP 7-1834022_A_T and CpG site cg12098896 among those without teen depression is in the opposite direction of the same association among those with depression. This interaction meQTL is of particular interest because both this SNP and CpG site are located within the gene *MAD1L1*, which has been previously found to contain SNPs associated with schizophrenia [103] and bipolar disorder [110], as well as a CpG site associated with major depressive disorder [20]. Moreover, the *MAD1L1* gene shows the highest transcript levels across tissues in EBV-transformed lymphocytes in the GTEx portal [43, 25]. EBV-transformed lymphocytes can show markers of depression [32] and stress [59] as a result of the process of transformation.

**Figure 7:**
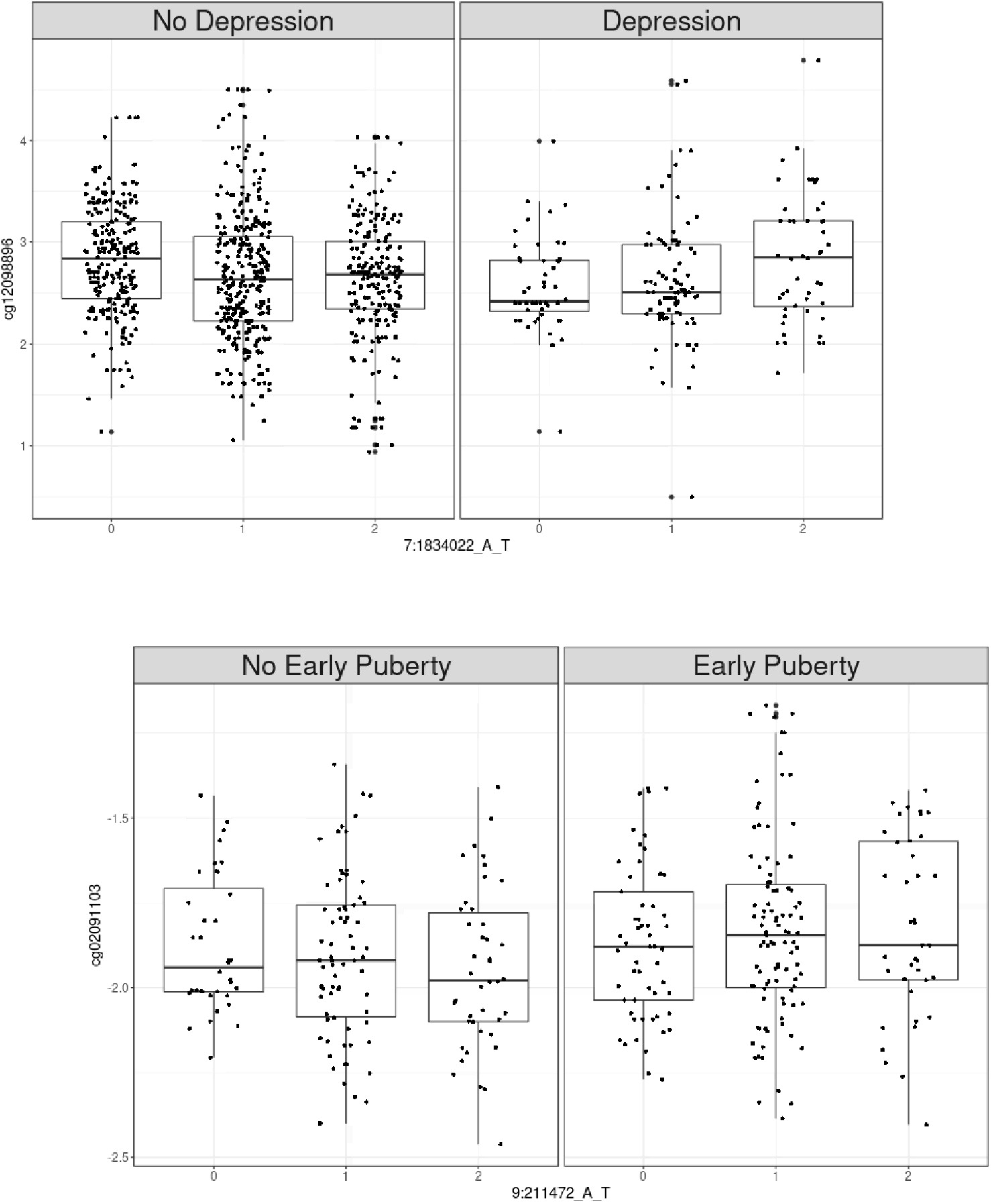
Example interaction meQTL boxplots. (a) An example of a depression-interaction meQTL with the SNP 7-1834022_A_T and CpG site cg12098896. For visual purposes, we stratify the depression scores for less than or equal to six versus depression scores greater than six. (b) An example of an early puberty-interaction meQTL with SNP 9-211472_A_T and CpG site cg02091103.

Another interesting depression-interaction meQTL contains a CpG site within the gene *CCNT2* (Table 5b), related to the cyclin family. It has been found to be associated with intellectual disability, particularly limitations in intellectual functions and social behavior [52]. In the GTEx portal, relative to other brain regions, this gene has high transcript levels in the cerebellar hemisphere and cerebellum regions of the brain, which play key roles in depression [55]. Our findings add to the known associations with these genes by suggesting the presence of a meQTL where the regulatory association differs by depression score.

For the puberty interaction meQTLs, an interesting puberty interaction meQTL targets the CpG site that is located within the *SPESP1* gene (Table 5a). This gene is related to breast cancer [27], as well as pre-eclampsia [107, 123]. Finally, it has also been found to be differentially expressed between male and female lung tissue [46]. Moreover, from the GTEx portal, this gene shows high levels of transcription in the testis and thyroid, tissues involved in puberty [101] and obesity [13], which is linked with early puberty. Our puberty interaction meQTLs suggest that regulation of these genes may be differential with respect to early puberty status.

We also performed enrichment analysis on the interaction meQTLs (Figure 8). For early puberty (Figure 8.a), we found four annotation categories (i.e., Coding, DNaseI digital genomic footprinting (DGF), Promoter Flanking, and Promoter) that were enriched for meQTLs (Bonferroni-corrected *p* < 0.05). In particular, among these four annotation categories, the DGF shows the greatest enrichment with 54.1% of the SNPs explaining 7.8% of early puberty-interaction meQTLs’ heritability. Coding regions have been found to be associated with Alzheimer’s disease [128]. In addition, promoter regions show enrichment, with 3.1% of SNPs co-localized in these regions explaining 13.1% of meQTL heritability. Promoter regions are known to drive cis-regulation of gene expression [43], and have been found to be associated with autism spectrum disorder [2] and breast and ovarian cancer [122]; a direct connection with early puberty is unclear.

**Figure 8:**
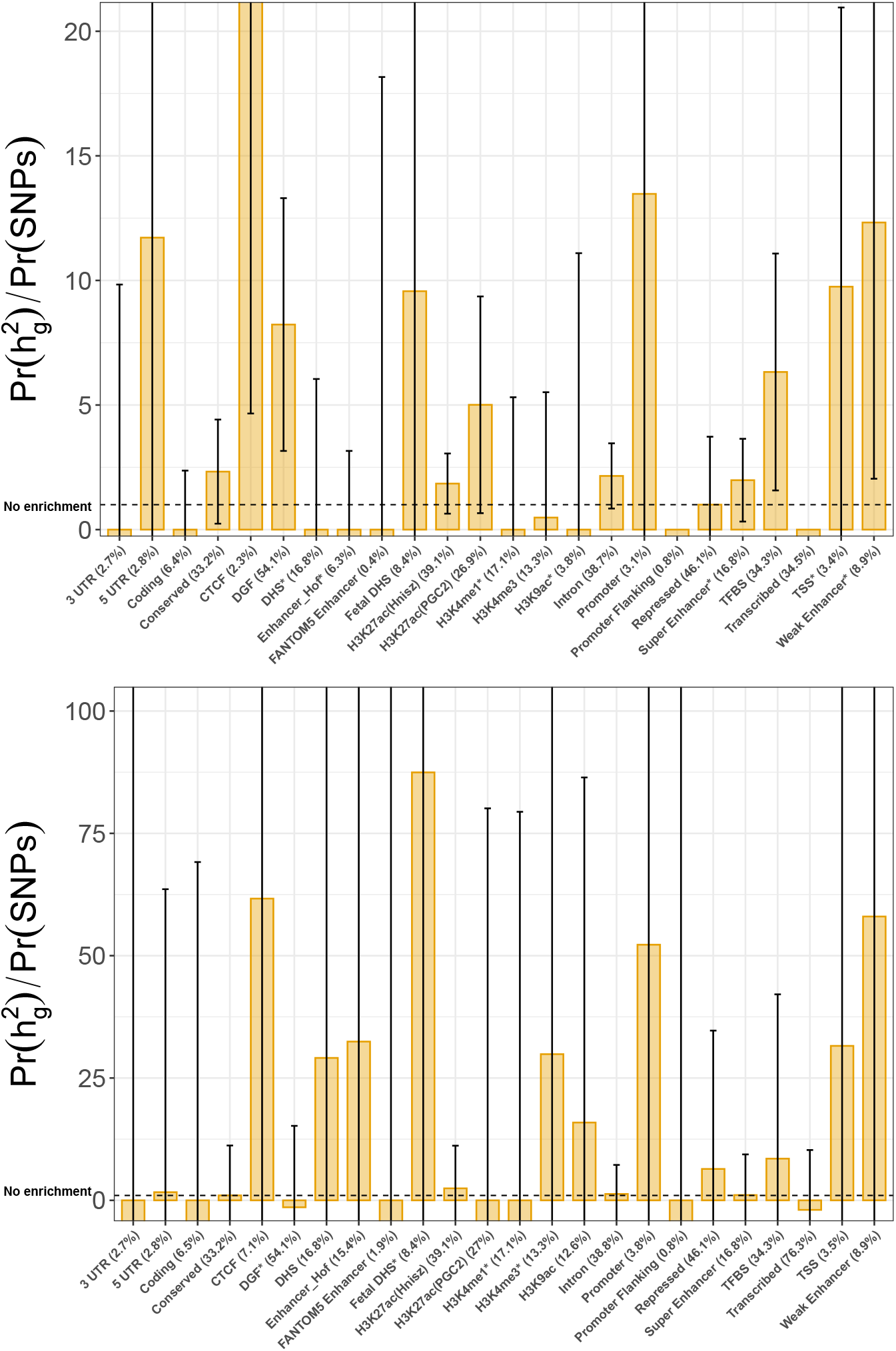
Enrichment analysis using stratified LD score regression for 24 annotation categories. Error bars show the jackknife standard errors around the estimates of enrichment, and the asterisk in the x-axis label represents significance (Bonferroni-corrected p < 0.05). (a) Enrichment analysis on the interaction meQTLs for early puberty. (b) Enrichment analysis on the interaction meQTLs for depression.

For the depression-interaction meQTLs, there are two annotation categories (i.e., DGF, and Weak Enhancer) that are enriched (Bonferroni-corrected *p* < 0.05; Figure 8.b). Among these two, the weak enhancer regions shows the largest enrichment estimate, with 8.9% of SNPs that are co-localized with weak enhancer regions explaining 58.2% of interaction meQTL heritability. The weak enhancer has been linked to mechanisms behind depression and suicide [21].

Although not significant, the the DNAse I hypersensitive site (DHS) markers show an enrichment with 16.8% of the SNPs explaining 89.9% of depression-interaction meQTLs’ heritability. The DHS has been found to be associated with obesity [83].

## Conclusions

In this work, we used genomic and phenotypic information in the Fragile Families Child Wellbeing Study cohort to study the interplay between DNA methylation and two traits that are important in adolescent health: teen depression and early puberty. We identified differentially methylated regions (DMRs), methylation quantitative trait loci (meQTLs), and interaction meQTLs across these data with respect to teen depression and early puberty status. This first genome-wide analysis of methylation data from the FFCWS cohort successfully characterized the impact of methylation on teen depression and early puberty. Because of the large sample size in this study and substantial representation of participants with early puberty or teen depression traits, we were able to show dynamic effects acting between ages 9 and 15, as well as the genetic regulation of methylation of CpG sites related to depression and early puberty status.

Our DMR analyses identified regions that were differentially methylated with respect to early puberty and teen depression status. We also found regions with differential methylation status across age with respect to both early puberty and teen depression. We identified 9,154,359 methylation QTLs and 51,191 interaction meQTLs with respect to early puberty status, and 70,569 interaction meQTLs with respect to depression score. We validated our meQTLs in the GoDMC Study data and also performed enrichment analyses with regulatory elements in the genome. Our findings have a broadly mechanistic interpretation into the biological underpinnings of early puberty and teen depression; many of the findings are potentially novel mechanistic genes that we will experimentally validate in the future [62]. Our analysis both recapitulates known biology and generates novel insights for understanding the mechanisms by which early puberty and teen depression manifest in a racially-diverse cohort, building the foundation for future interventions to improve adolescent health.

## Competing interests

BEE is on the SAB of Celsius Therapeutics, Freenome, and Creyon Bio. BEE was employed by Genomics plc from July 2019-August 2020.

## Author’s contributions

RDV, ING, and BEE conceived the experiments. LMS performed data collection and some quality control. RDV, ING performed quality control on the methylation data. DA performed quality control and imputed the genotype data. RDV and ING built the methods and ran the experiments. AV assisted with functional validation of the results. RDV, ING, and BEE analyzed the results. RDV, ING, and BEE wrote the paper. SM initiated the FFCWS, ran the study for over twenty years, and supported this work. DAN and CM designed and funded FFCWS genotyping and methylation studies up to the point of data collection.

## Acknowledgements

BEE was supported by NIH R01 HL133218, a Sloan Faculty Fellowship, and NSF CAREER AWD1005627. Research reported in this publication was supported by the Eunice Kennedy Shriver National Institute of Child Health and Human Development (NICHD) of the National Institutes of Health under award numbers R01HD36916, R01HD39135, and R01HD40421, as well as a consortium of private foundations. Research reported in this publication was also supported by the National Library Of Medicine of the National Institutes of Health under Award Number T32LM012411. DNA data collection was funded under NIH grants R01 HD076592 and R01 HD073352. Methylation data collection was funded by NIH grants R01 MD011716 and R01 MH103761. The content is solely the responsibility of the authors and does not necessarily represent the official views of the National Institutes of Health.

## Notes

https://fragilefamilies.princeton.edu/

