## Supplemental Information for "Differentially methylated regions and methylation QTLs for teen depression and early puberty in the Fragile Families Child Wellbeing Study"

### Supplementary Materials for "Differentially methylated regions and methylation quantitative trait loci for depression and early puberty in the Fragile Families Study "

Roberta De Vito*, Isabella N. Grabski*, Derek Aguiar, Lisa M. Schneper, Archit Verma, Juan Castillo Fernandez, Jordana Bell, Sara McLanahan, Daniel A. Notterman, and Barbara E. Engelhardt

#### A. Methods

##### A.1 Study design and subjects phenotyping

As we described in the main manuscript, for adolescent depression, the measured phenotype is a continuous score based on the survey data at year 15, with a scale from 0 to 16.

The score is based on 5 variables (Table 1, main manuscript) defined by the Center for Epidemiologic Studies Depression Scale (CES-D) and adopted in the National Longitudinal Study of Adolescent Health, Wave I. The CES-D items is based on the person's feelings using values on a four-point ranging from 1(strongly agree) to 4 (strongly disagree). In the Fragile family Data in Year 15, items are specified from the past four weeks.

For early puberty, we consider when the child's breasts begun to grow. The answer is divided in three responses (i.e. No, Yes, barely,Yes, definitely). the exact question for the Mother's was "Would you say that her breasts have started to grow? Would you say..". As we stated in the manuscript, we transform this variable in a dichotomous replies (i.e., No and Yes).

##### A.2 Differences between M-values and Beta-values

The Beta-value is the ratio between the methylated probe intensity and the overall intensity. Following the notation used by Illumina, we have for each CpG site:

$$Beta_i = \frac{\max y_{i,methyl,0}}{\max y_{i,unmethyl,0} + \max y_{i,methyl,0} + \alpha}, \tag{1}$$

*Equal contributor

where $y_{i,methyl}$, $y_{i,unmethyl}$ are the intensities measured by the $i^{th}$ methylated and unmethylated) probes respectively. Negative values are set to zero to circumvent negative values. Also, Illumina suggests to add the constant $\alpha$, generally set to 100, to regularize the beta value. Beta-values are between zero and one, or zero and 100%.

The M-value is computed as:

$$M_i = \log_2 \left( \frac{\max y_{i,methy}, 0 + \alpha}{\max y_{i,unmethy}, 0 + \max y_{i,methy}, 0 + \alpha} \right). \quad (2)$$

When the M-value is close to zero, the intensity between the methylated and unmethylated signal is similar, resulting in a CpG site that is half-methylated. When the M-value is positive or negative, the probes is methylated or unmethylated respectively.

In a paper by Du et al. (2010), a comparison between the two values is provided. Firstly they provide the relationship between the Beta-value and M-value:

$$Beta_i = \frac{2^{M_i}}{2^{M_i} + 1} \quad M_i = \log_2 \left( \frac{Beta_i}{1 - Beta_i} \right). \quad (3)$$

They proceed with a performance comparison between the Beta and M-values, highlighting the differences between these two. In fact, even if the Beta-value can be interpreted as the percentage of a site that is methylated, it has several statistical issues, such as heteroscedasticity outside the middle methylation range, with serious consequences for applying many statistical models. The M-value is approximately homoscedastic, allowing a wider range of statistical methods in downstream analysis that require homoskedasticity.

### B Results

#### B.1 Validation in the GoDMC Study

The GoDMC study identified meQTLs calculated in whole blood DNA samples with HumanMethylation450 or EPIC array profiles from 32,851 participants in the GoDMC cohort Min et al. (2020). In the manuscript we have reported some plots that show an enrichment between the two data sets. Here we also report the q-q plot (Figure S1) of the observed p-values for the meQTLs in common between the two data-sets compared with the expected p-values. We see a strong enrichment of low p-values as compared to the distribution of p-values expected by chance (Figure S1). This validation further emphasizes that the meQTLs identified in the FFCWS replicate in an outside sample, thus suggesting that the meQTLs represent general biological signal.

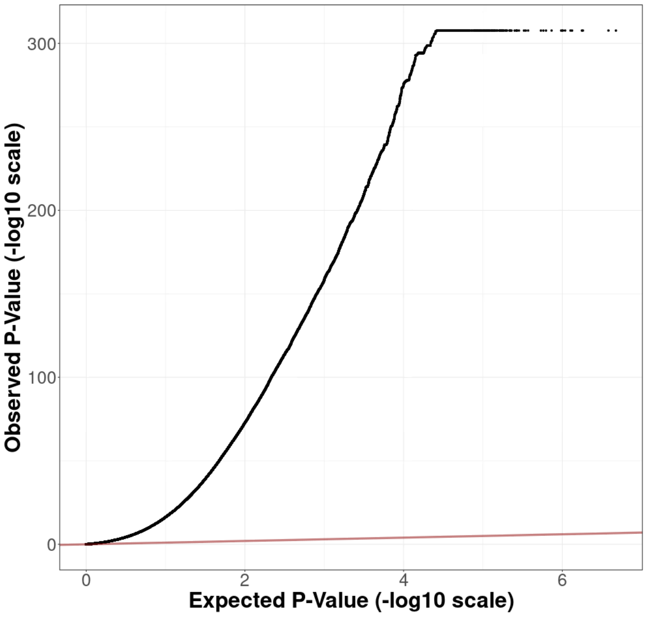


Figure S1: Quantile-quantile plot of observed and expected p-values. Quantileâquantile plot (QQ-plot) of ranked expected (x-axis) versus observed (y-axis) p-values for the meQTLs found in common between our work and the GoDMC study. The straight line (in dark red) represents the identity ($x = y$) line.

Min, J. L., Hemani, G., Hannon, E., Dekkers, K. F., Castillo-Fernandez, J., Luijk, R., Carnero-Montoro, E., Lawson, D. J., Burrows, K., Suderman, M., et al. (2020). Genomic and phenomic insights from an atlas of genetic effects on DNA methylation. *medRxiv*.
